# Simulating the Impacts of Climate Change on UH Mānoa Lettuce (*Lactuca sativa*) Growth by Modifying Air Temperature, Soil Water Availability, and Atmospheric CO_2_ Concentration

**DOI:** 10.1101/2025.08.13.670242

**Authors:** Nicholas Yos, Camilo Mora, Kira Webster, Kyle McDowell

## Abstract

Plant species are adapted to survive under specific ranges of temperature, water availability, and atmospheric CO_2_ concentration. Climate change-induced shifts in these environmental conditions have the potential to significantly affect nearly all terrestrial plants. A number of studies have explored the impacts of changing one or two of the conditions listed above, but few have examined the combined effects of all three. To study the cumulative influences of the three environmental conditions, 350 Mānoa lettuce (*Lactuca sativa*) plants were grown in indoor growth chambers. Within the chambers, plants were grown under varying degrees of CO_2_ concentration, water availability, and temperature for 21 days. At the end of this period, the leaf mass (biomass) of each plant was cut, dried, and weighed. Percent mortality and nitrogen content were also measured. Across all combinations of temperature and water availability, elevated CO_2_ concentrations were associated with increased biomass production and survival rates. Survival and biomass decreased under high temperatures and both high and low water availability. The combination of environmental conditions that produced the largest amount of biomass was 750 ppm CO_2_, 80% of soil water capacity, and 24 °C while the treatment that produced the least amount was ambient CO_2_, 60% of soil water capacity, and 36 °C. Nitrogen content increased under high temperatures and water availability. These results suggest that although increased atmospheric CO_2_ levels have the potential to promote lettuce growth, lettuce yield is still likely to decrease in many regions due to the negative effects of high temperatures, drought, and flooding.

## 1.0 INTRODUCTION

### 1.1 BACKGROUND

The terrestrial biosphere and human civilization are fundamentally dependent on plants. As the dominant primary producers in nearly all terrestrial environments, plants provide the biomass and energy that fuel terrestrial ecosystems, which store approximately 2,300 gigatons of carbon in the form of vegetation and soil [1]. The roots of grasses and trees mitigate erosion on both slopes and the coast [2] and can improve water capture and quality [3–4]. Plant based agriculture directly provides approximately 82% of the global calorie supply [5], takes up half of the world’s habitable land area through plant farming and plant-dependent animal agriculture [5], and produces a total global value of well over four trillion US dollars a year [5]. Global forest products are worth approximately 244 billion US dollars a year [5] and 81% of the 450,000 multifamily units built in the United States in 2023 were wood-framed [6]. Without healthy, stable plant communities, terrestrial biomes and human societies cannot exist.

### 1.2 EFFECTS OF CLIMATE CHANGE ON PLANT GROWTH

Due to the global importance of plants, any threats to plant life posed by climate change have the potential to lead to severe global consequences. All organisms are evolved to survive within certain ranges of environmental conditions and changes to these conditions can negatively impact the ability of an organism to survive within an environment [7]. Plants require water, atmospheric CO_2_, and a favorable temperature regime to survive, all of which can be significantly altered by the effects of climate change [7]. This study explores the individual and collaborative impacts of all three of these factors on the survival, growth, and nutritional quality of lettuce.

#### 1.2.1 IMPACTS OF HIGH TEMPERATURES

High temperatures present a threat to a number of plant functions, including photosynthesis, the production of biomass and reactive oxygen species (ROS), and reproduction. During extreme heat events, plants are known to experience damage to chloroplast protein complexes, chlorophyll, and grana stacks, all of which are vital to the normal functioning of photosynthesis [8–11]. This is compounded by alterations in the activity of vital enzymes such as Rubisco and chlorophyllase [8–9]. Due largely to this decrease in photosynthesis, many plant species have been observed to exhibit reduced rates of growth and biomass production under high heat conditions [8]. This reduction can lead to decreased yield in important food crops such as wheat, rice, and maize [10]. Heat-induced damage to enzymes and chlorophyll can also lead to the accumulation of dangerous ROS, such as superoxide, hydrogen peroxide, and hydroxyl radicals [8–10]. The presence of these molecules inside cells can lead to the irreversible oxidation of proteins, DNA, and fatty acids [12]. During periods of flowering and seeding, high temperatures can lead to poor seed production or sterility due to disturbances in reproductive organ development, the fertilization process, and embryo growth [8, 10, 13]. These disturbances are reflected in the seedlings of heat-impacted parent plants, which experience higher levels of mortality and lower growth rates [10]. All of these factors contribute to reduced plant success, which can lead to rippling effects on ecosystems and human communities.

#### 1.2.2 IMPACTS OF LIMITED WATER AVAILABILITY

Water stress due to drought can lead to many of the same impacts on plants as extreme heat, but for differing reasons. Leaf stomata often close under drought conditions to reduce evapotranspiration, but this also reduces CO_2_ uptake and, by extension, the amount of CO_2_ that is available for photosynthesis [14–16]. Low water availability also limits photosynthesis through chloroplast disfiguration, pigment synthesis reduction, disruptions to the thylakoid electron transport chain and Calvin cycle and, similarly to heat stress, a general reduction in the effectiveness of photosynthetic enzymes [14–16]. Many of these limitations can be attributed to the actions of ROS, which can also be created due to water stress-induced NADPH accumulation [9, 14]. As well as reducing carbon assimilation through impaired photosynthesis, water deprivation can limit plant growth by preventing water flow to new cells and inhibiting mitosis [14, 17]. This can lead to reductions in root and shoot development, leaf size, crop yield, and reproductive success [9, 14, 17]. Extended exposure to drought conditions has the potential to cause widespread leaf surface area reduction, leaf loss, air bubble formation in xylem tissue, a complete cessation of photosynthesis, significant metabolism disruption, and eventually plant death [9, 14–15].

#### 1.2.3 IMPACTS OF ELEVATED CO_2_ CONCENTRATIONS

Increased atmospheric CO_2_ concentration has the potential to mitigate some of the negative effects of water and temperature stress, but only to an extent. Exposure to CO_2_-enriched air has been found to stimulate increased photosynthesis in a wide variety of plant species, which promotes higher levels of growth and biomass production [18]. This increase in primary production and carbon assimilation has the potential to slightly reduce the rate of rising CO_2_ concentrations, although it is unlikely to be significant enough to provide any substantial mitigation of climate change [18–19]. Additionally, increased CO_2_ concentrations have been found to promote water use efficiency (WUE), which refers to the amount of water that is lost through evaporation for every unit of CO_2_ assimilated [13, 18–19]. Increased WUE could allow plants to survive regional drying trends and exhibit higher levels of primary production under low water conditions [18–19]. However, while improved WUE and photosynthetic capacity are clearly visible under ideal laboratory conditions, these improvements become less significant when other environmental factors such as competing species, nutrient limitation, and the soil microbiome are introduced [20–22]. Additionally, the effects of increased CO_2_ have been found to decrease over time as plants gradually acclimate to higher concentrations [19, 22–23].

#### 1.2.4 COMBINED IMPACTS

It is important to study temperature, water availability, and CO_2_ concentration simultaneously because these three factors can have interacting impacts on the health of a plant. One clear example of this can be seen in the case of stomatal conductance. The stomatal conductance of a plant is a measure of the exchange of gases like CO_2_ and water vapor through the stomata and is primarily controlled by the degree to which leaf stomata are opened or closed [24]. Increased atmospheric CO_2_ concentrations allow plants to obtain CO_2_ with lower stomatal conductance and water loss, which is why increased CO_2_ levels tend to improve plant WUE [13, 18–19, 25]. Alternatively, stomatal closure due to severe drought conditions could partially negate any stimulatory effects of increased CO_2_ [14, 25]. The problem of stomatal conductance can be complicated further by high temperatures, which can trigger stomatal closure or opening regardless of CO_2_ or water availability [9, 26]. The impacts on photosynthesis and metabolism caused by any of the three can also play a role in the control of stomata and other plant functions [13]. Complex biological interactions such as those that control stomatal conductance can be found in most species and can play an important role in the survival of an organism [13]. However, many studies into the potential impacts of climate change do not take these complexities into consideration [13].

### 1.3 EFFECTS OF CLIMATE CHANGE ON PLANT NITROGEN CONTENT

Aside from the effects of climate change on plant growth, changing climatic conditions - especially increases in atmospheric CO_2_ concentration - have the potential to negatively impact the protein content of crop plants. While it is true that increased CO_2_ concentrations can encourage many crop plants to grow more quickly, numerous studies have found that food produced under elevated CO_2_ levels often has lower levels of protein [27–30] and nutrients [27, 30] when compared to crops grown under lower atmospheric CO_2_. Multiple possible explanations have been proposed for the phenomenon of reduced plant protein content under high CO_2_ concentration. One potential cause is the biomass dilution effect that occurs when plants produce a disproportionately large quantity of nonstructural carbohydrates due to atmospheric CO_2_ enrichment, which can result in crops that have a lower nitrogen content per unit of biomass [31–34]. Increased photosynthetic efficiency at elevated CO_2_ concentrations can also increase photosynthetic nitrogen use efficiency by decreasing the need for photosynthetic enzymes and increasing carboxylation efficiency, thus reducing plant nitrogen demand and eventually crop nitrogen content [33, 35]. A third factor that can influence the nitrogen content of plant matter is decreased stomatal conductance at high atmospheric CO_2_ concentrations, which reduces transpiration and slows down the uptake of nitrogen from roots [31, 33, 36]. Regardless of the cause or combination of causes that lead to lower nitrogen concentrations in crops grown under elevated CO_2_ concentrations, the decreased nutritional value of the food produced has the potential to partially negate the beneficial effects of faster growth rates.

### 1.4 LETTUCE AS A MODEL SPECIES

Lettuce was selected as a model organism to explore the impacts of climate change on plant growth because of its global economic importance, small size, fast growth rate, and sensitivity to changing environmental conditions. Lettuce is widely grown on both industrial and small farms around the world and is a relatively inexpensive and nutritious food source [37]. Over 26.8 million tons of lettuce were grown in 2017 [37] and the total value of the 2020 global crop of lettuce and the related species chicory was approximately three billion US dollars [5]. Lettuce is useful as a model organism for studying climate change because it is a relatively small, fast-growing plant that has been shown to be particularly sensitive to changes in temperature and water availability [38–39]. Any findings collected on lettuce could potentially be extended to other members of the Asteraceae family, which includes sunflowers, safflowers, chicory, and artichokes among at least 32,000 other species. However, it should be noted that different plant species can respond to climate change in very different ways and the response of lettuce to climate variation should not be expected to be exactly mirrored in other species [22].

### 1.5 HYPOTHESIS

Our primary hypothesis for this study was that lettuce growth would be negatively impacted by both increasing temperatures and decreasing water availability, but that elevated CO_2_ concentrations have the potential to reduce the negative impacts caused by these two environmental factors. Additionally, we hypothesized that the lettuce biomass produced under elevated CO_2_ concentrations would have a lower nitrogen content than lettuce grown under ambient concentrations.

## 2.0 METHODS

### 2.1 MORA LAB GROWTH CHAMBERS

This experiment utilized the Mora Lab Intelligent Plant Growing System (IPS) growth chambers (see Fig 1), which are located in the UH Mānoa Department of Geography and Environment [40]. Each of the ten chambers consists of a 2 m by 1 m by 1 m metal frame enclosed by reflective insulating material and each is capable of containing up to 40 two-liter plant pots in five rows that are suspended approximately 0.2 m above the bottom of the chamber. Lighting within the chambers is provided by 8 Unifun grow lights located approximately 0.5 m above the top of the pots. The temperature, soil moisture, and CO_2_ availability within each chamber can be independently regulated, allowing researchers to study the effects of changing climate conditions on plant growth.

**Fig 1.**
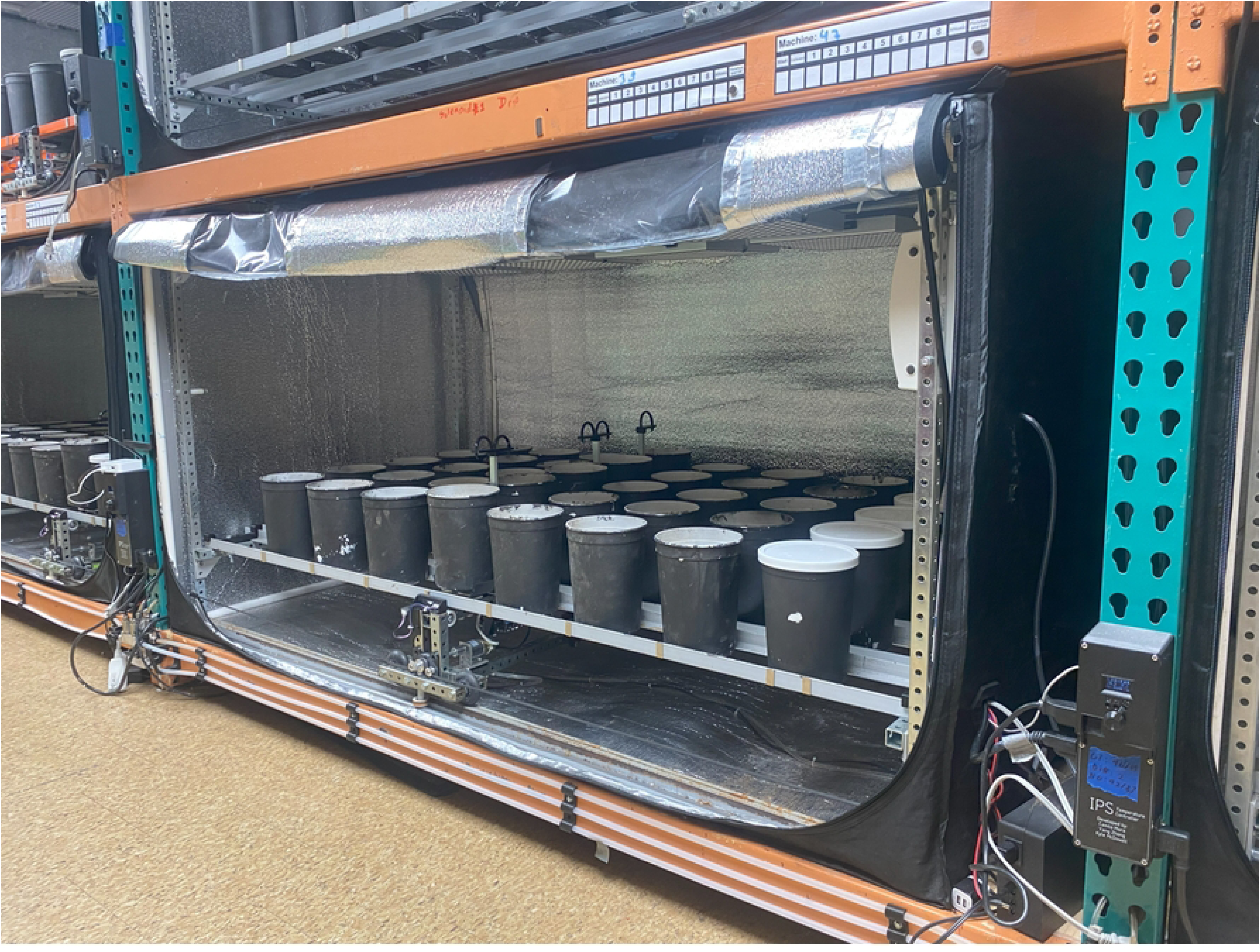
One of the Mora Lab IPS growth chambers with the door open (picture by author)

#### 2.1.1 TEMPERATURE REGULATION

The temperatures of the chambers are regulated by custom-made IPS temperature controllers (Fig 2) [41]. Each of these devices controls a small heater and air conditioner contained within the chambers as well as the day-night light cycle. When temperatures inside the chamber are below the set temperature - as determined by a built-in PT100 probe - the controller turns on the heater for short bursts at regular intervals until temperatures reach the desired level. When temperatures are too high, the controller turns the air conditioner on steadily until chamber temperatures match the set temperature. To ensure a consistent temperature throughout the chamber, two fans provide constant air circulation.

**Fig 2.**
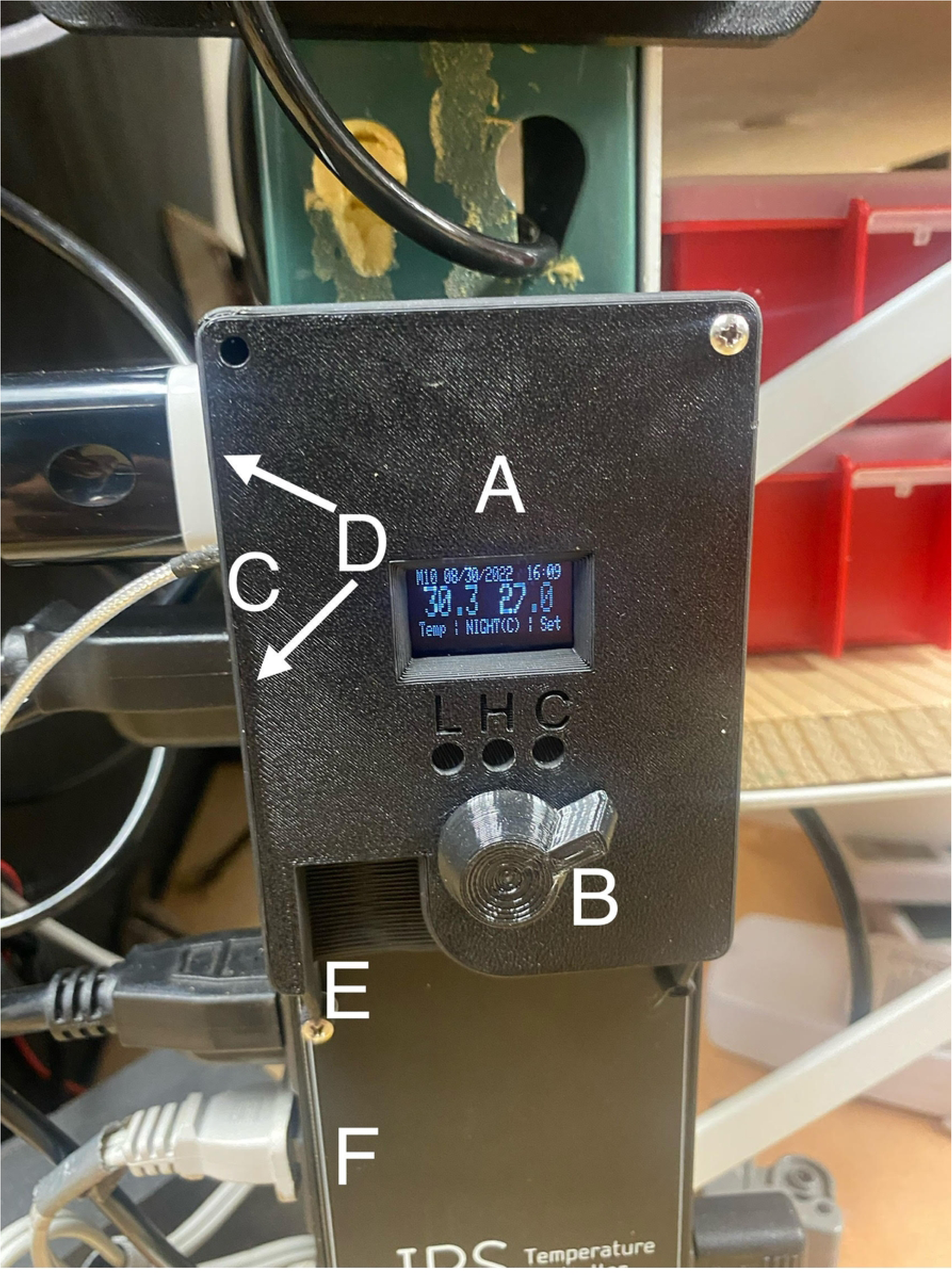
Picture of the IPS temperature controller with key features labeled (picture by author) The controller allows the operator to regulate the temperature and day-night light cycle of each chamber, shown on the display (A) and controlled by a dial (B). Temperature is measured by a PT100 temperature probe (C). The top two outlets (D) on the side of the controller provide 110V AC power, the third outlet (E) turns the grow lights on and off based on the programmed day-night cycle, and the bottom two outlets (F) power an air conditioning unit and heater than turn on and off based on the internal temperature of the chamber.

#### 2.1.2 WATERING GANTRY

The amount of water provided to the plants and thus the soil moisture within each pot was based on the field capacity of the growth medium. The field capacity of a soil is the maximum amount of water it can contain without the effects of gravity causing any of the water to drain downwards [42]. For this experiment, the field capacity of the media used - coarse vermiculite - was approximately determined by saturating a known weight of media in a pot with drainage holes in the bottom, covering the top of the pot to prevent evaporation, and re-weighing the pot after three days. The initial and final weights were used to determine the weight of water that could be contained by a given volume of media. By determining the field capacity, the amount of water provided could be expressed as a fraction of the maximum weight of water the medium could contain.

The quantity of water added to each pot was controlled by the IPS robotic watering gantry [43]. This machine consists of a motorized rolling carriage that contains five load cells that control five water spigots, one for each row of pots. When the machine is switched on, it rolls parallel to the rows of pots until it detects the presence of a magnet, which are placed at the front of each column of pots. Upon stopping, the machine rises upwards and the five load cells determine the weight of each pot in the column. If the weight of a pot is below the programmed level, the spigot releases water into the pot until it reaches the desired weight. This process is repeated until every column has been weighed and watered, at which point the gantry returns to the starting point. See Fig 3. for an image of the gantry in a lifted position.

**Fig 3.**
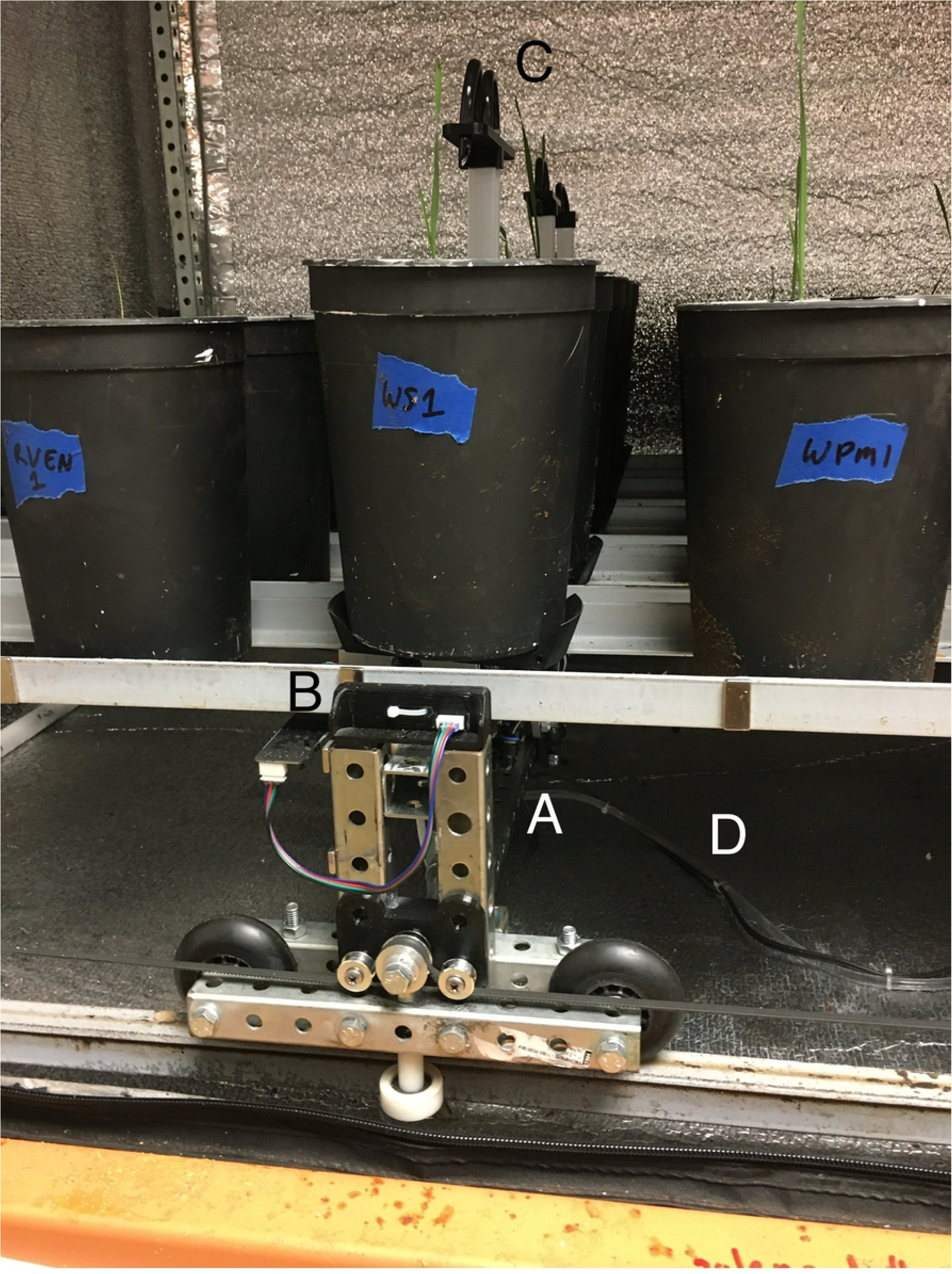
Picture of watering gantry with a row of pots lifted for weighing and watering (picture by author) When turned on, the motorized gantry (A) rolls along the track until the magnetic sensor (B) detects the presence of a magnet. The gantry then raises and weighs each pot individually. If the weight of a pot is below the set weight, the five spigots (C) add water to each pot until they reach the set weight. The gantry then lowers and moves on until it detects the next magnet. Water is provided by a tube (D) connected to an outside water source.

**Fig 4.**
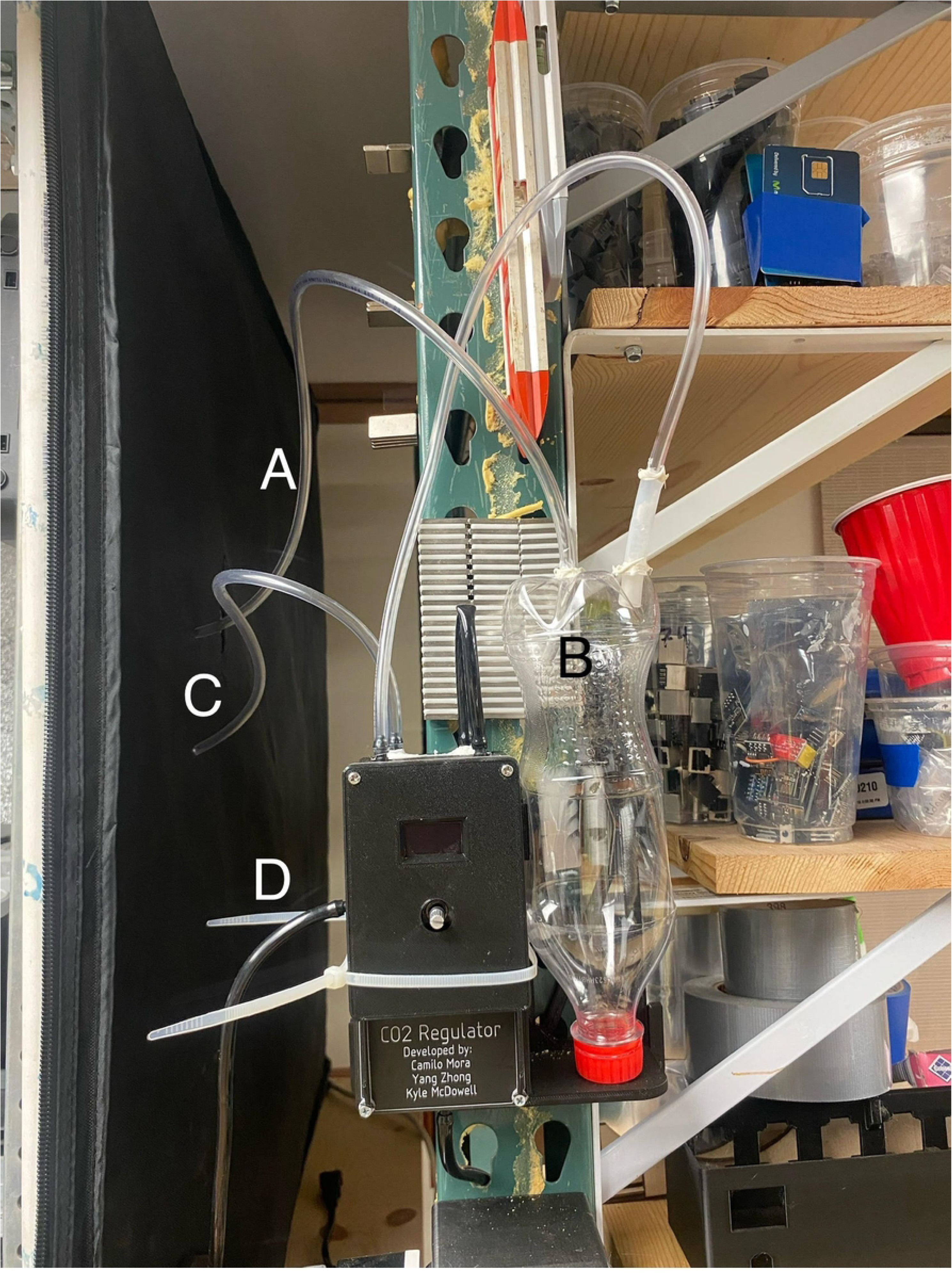
Picture of the IPS CO_2_ regulator (picture by author) Tube A draws in air from the chamber to measure CO_2_ concentration. A plastic bottle (B) is used to capture excess moisture before it reaches the device. If CO_2_ concentrations are below the set level, a pulse of pure CO_2_ is added via tube C. This CO_2_ comes from tube D, which is attached to an external CO_2_ canister. (Picture by author)

#### 2.1.3. CO_2_ REGULATION

Elevated atmospheric CO_2_ concentrations were simulated by the addition of pure CO_2_ gas from an external gas canister. The addition was controlled by a custom CO_2_ regulator designed by Camilo Mora, Kyle McDowell, and Yang Zhong. These devices constantly measure the CO_2_ concentrations inside each chamber and add a short burst of gas whenever concentrations drop below the desired level. In the chambers without elevated CO_2_ levels, a corner of the door of the chamber was left slightly open to allow the air in the chamber to mix with the external atmosphere. Because the ambient CO_2_ concentration was influenced by the number of people in the building, CO_2_ levels in the ambient chambers ranged from 430-520 ppm daily.

### 2.2 LETTUCE GROWTH PROCEDURES

For this experiment, Mānoa lettuce (*Lactuca sativa*) plants were grown under a range of potential climate change conditions. Lettuce seeds - purchased from the UH Mānoa CTAHR seed project - were sprouted in SkinnyBunny rock wool cubes which were then transplanted to pots of ThermoRock coarse vermiculite in the growth chambers at the 2-4 leaf stage. All plants were approximately the same size and age upon planting. Before planting, ten grams of Island Supreme 16:16:16 NPK fertilizer was mixed in with the top three centimeters of vermiculite in each pot. After allowing the seedlings to acclimate to the pots for one week under the range of soil moisture levels that would be used in the experiment, the chambers were closed and the temperature and CO_2_ conditions in each were set. Of the ten chambers, half were left at ambient CO_2_ concentration, which ranged from 430-520 ppm daily. CO_2_ concentrations in the other five chambers were elevated to 700 ppm, meant to represent global CO_2_ concentrations in 2100 under moderate emissions (SSP2-4.5) [44]. Within each group of five chambers, the daytime temperatures of the chambers were set at four-degree intervals from 20° C to 36° C. At night, temperatures inside each chamber were decreased by five degrees. Within each chamber, each row of pots was given a different degree of soil moisture, ranging from 100% of field capacity to 60% of field capacity. Machines were watered once a day for the duration of the experiment. Grow lights were turned on for 12 hours a day to simulate a day-night cycle. Once these conditions were established, the lettuce plants were allowed to grow for 21 days before harvesting. See Fig. 5 for an image of the interior of a growth chamber during the growing period.

**Fig 5.**
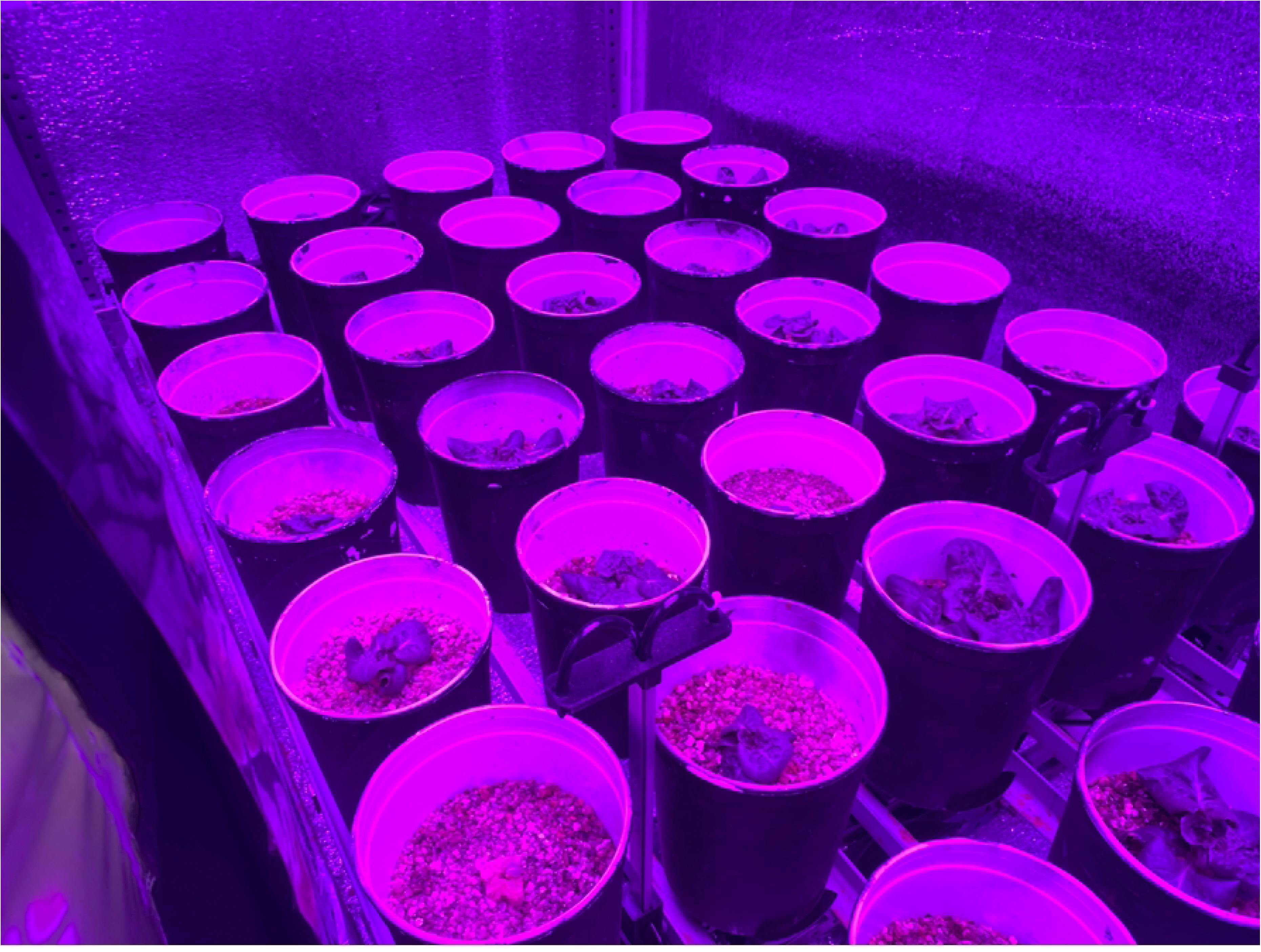
Lettuce growing in the chambers.

### 2.3 DATA COLLECTION

The three variables that were recorded in this experiment were plant mortality, leaf dry biomass, and leaf nitrogen content. Throughout the experiment, plant mortality across all treatments was recorded regularly as determined by visual inspection. After 21 days, each plant was cut at the base and all aboveground biomass was collected, dried for 24 hours at 70 °C in a Fisher Scientific Isotemp Oven, and weighed with a Ohaus Adventurer scientific scale. Following this process, three random samples from different plants within each experimental group (each unique combination of temperature, water availability, and CO_2_ treatments) were sent to the UH Mānoa Agricultural Research center to be tested for nitrogen content, which can be used as a proxy for protein content. For experimental groups that had 3 or less living plants remaining by the end of the experiment, all remaining plants were tested. After data collection was complete, the information we collected on mortality, leaf mass, and protein content was processed via regression analysis in R.

### 2.4 DATA ANALYSIS

All data collected in this study was analyzed using code designed by [40]. To adequately represent the nonlinear relationships between the different independent variables examined in this experiment, this code used regression analysis to describe the interactions of the three environmental factors studied. To determine the best model for this analysis, several possible models were compared and the one that produced the smallest AIC value while also containing both temperature and water was selected to process the data. Of the six models used in the regression analysis for this study, five incorporated both first and second order polynomials. Lower order components were included in any models that featured interaction terms. See S1 Table for the models used in the data analysis. Figures 6 through 11 were created in RStudio 4.3.1 by plotting the raw data produced by the regression analysis on contour plots with subplots that represent the individual interactions between pairs of variables.

**Fig 6.**
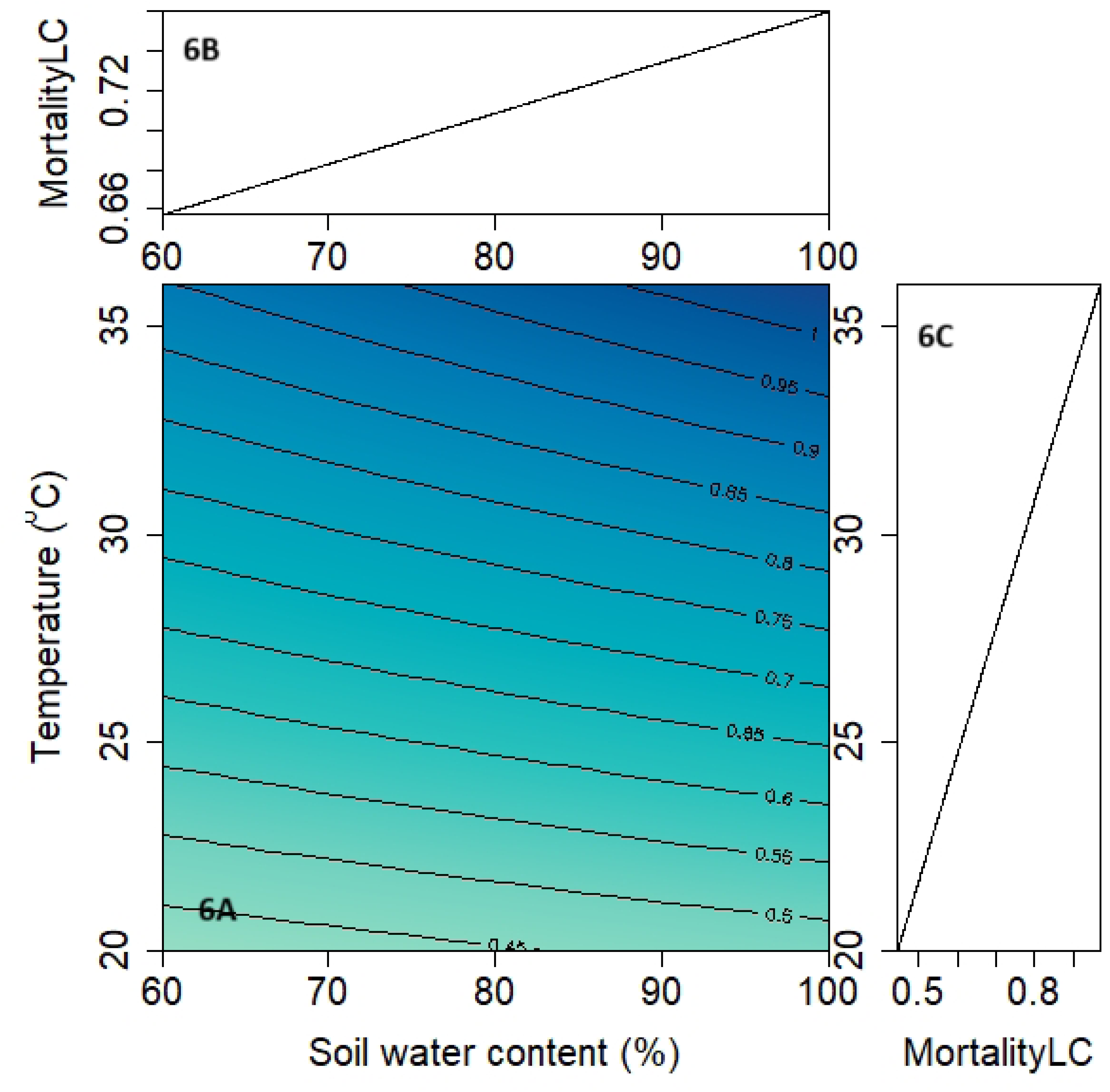
Impacts of temperature and water availability on lettuce mortality under ambient (LC) and elevated (HC) CO_2_ concentrations. The central figures (6A) show the combined impacts of temperature and water availability on mortality, with darker colors representing higher survival rates. See contour lines for specific mortality rates, which range from 0 for complete mortality to 1 for complete survival. The upper subplots (6B) show the relationship between mortality and water availability alone and the lower subplots (6C) show the relationship between mortality and temperature alone.

## 3.0 RESULTS

### 3.1 STATISTICAL ANALYSES

Regression analysis found that, although most predictor variables used in this experiment were not statistically significant, the regression models fit the data in five out of six models in S1 Table. The model comparing the effects of water availability and temperature on leaf mass under elevated CO_2_ concentrations was the only model in which either of the predictor variables had p-values under 0.05. In all other models, the p-values for the t-statistics of the predictor variables ranged from 0.05 to 1, indicating a lack of statistical significance for individual variables. See S2 Table for specific p-values for predictor variables. With this being said, the p-values for the F-statistics of nearly every regression model were well below 0.05, the only exception being the model examining nitrogen content under elevated CO_2_. These p-values indicate that, as a whole, five of the six models were generally accurate representations of the expected values of the response variables.

### 3.2 MORTALITY

The effects of the environmental conditions studied in this experiment on plant mortality are compiled in Figs 6 and 7.

**Fig 7.**
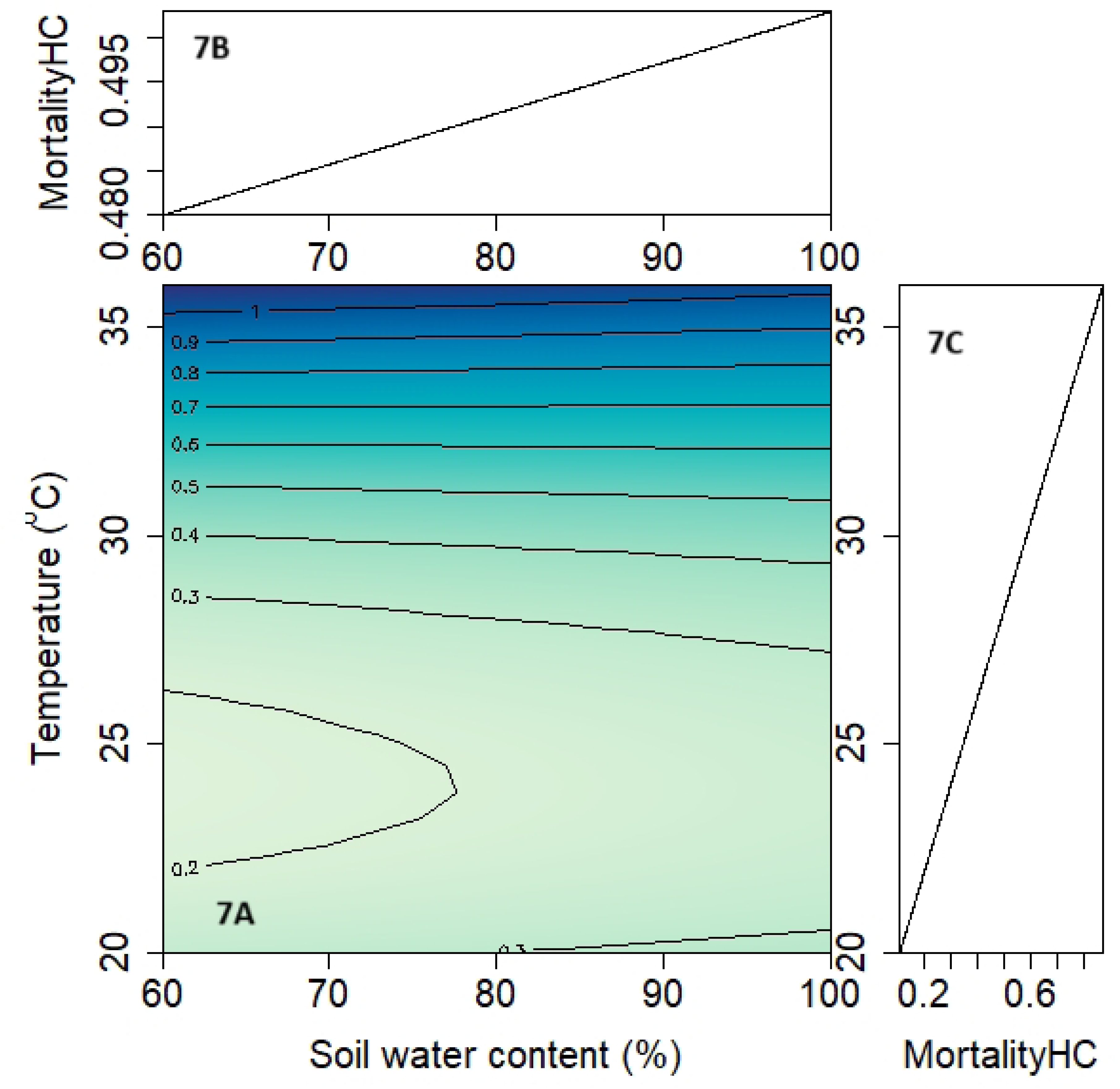
Impacts of temperature and water availability on lettuce mortality under ambient (LC) and elevated (HC) CO_2_ concentrations. The central figures (7A) show the combined impacts of temperature and water availability on mortality, with darker colors representing higher survival rates. See contour lines for specific mortality rates, which range from 0 for complete mortality to 1 for complete survival. The upper subplots (7B) show the relationship between mortality and water availability alone and the lower subplots (7C) show the relationship between mortality and temperature alone.

As seen in Fig 6, the highest survival rates were found under low temperatures and low water availability. Mortality increases at a relatively linear rate as both temperature and water availability increase. In Fig 7, mortality also increases at a linear rate with increasing temperatures, but soil water availability has a very limited impact on mortality. Overall, mortality was lower under elevated CO_2_ concentrations than under ambient conditions.

#### 3.2.1 RELATIONSHIP WITH TEMPERATURE

High mortality rates occurred across most chambers used in this experiment, with mortality generally increasing with temperature. However, neither the ambient nor elevated CO_2_ chambers experienced a consistent increase in mortality with increasing temperatures. For the ambient CO_2_ chambers, the coldest chamber experienced the lowest mortality (34%), yet the second coldest chamber was tied for the second highest mortality (89%). For the elevated CO_2_ chambers, the lowest mortality rate (14%) occurred in the second coldest chamber, with the coldest chamber having the third highest mortality rate (34%). Both the ambient and the elevated chambers set at 36 °C experienced 100% mortality, although it took the elevated chamber 14 days longer to reach complete mortality. See Table 1 and S1 and S2 Figs for a more complete comparison of temperature and mortality at the end of the experiment and over time.

**Table 1:** Final percent mortality of lettuce plants at ambient or elevated CO_2_ concentrations under different temperature regimes.

| Daytime temperature (°C) | Final percent mortality at ambient CO<sub>2</sub> (%) | Final percent mortality at elevated CO<sub>2</sub> (%) | Increase in survival between ambient and elevated CO<sub>2</sub> (%) |
| --- | --- | --- | --- |
| 36 | 100 | 100 | 0 |
| 32 | 89 | 74 | 125 |
| 28 | 43 | 23 | 35 |
| 24 | 89 | 14 | 650 |
| 20 | 34 | 34 | 0 |

#### 3.2.2 RELATIONSHIP WITH SOIL WATER CONTENT

Soil water content had a very small effect on lettuce mortality in the elevated CO_2_ chambers, with a <2% increase in mortality as water availability increased from 60% to 100%. This effect was more pronounced in the ambient CO_2_ chambers, where plants experienced an 18% increase in mortality as water availability increased. See Table 2 and S3 and S4 Figs for a comparison of mortality over time for plants grown under different soil water content levels.

**Table 2:** Percent mortality of lettuce plants at ambient or elevated CO_2_ concentrations under different water availability regimes.

| Soil water availability | Final percent mortality at ambient CO<sub>2</sub> (%) | Final percent mortality at elevated CO<sub>2</sub> (%) |
| --- | --- | --- |
| 100% | 83 | 49 |
| 90% | 71 | 51 |
| 80% | 57 | 31 |
| 70% | 71 | 51 |
| 60% | 63 | 43 |

#### 3.2.3 RELATIONSHIP WITH CO_2_ CONCENTRATION

Chambers with elevated CO_2_ generally experienced lower mortality rates than those with ambient CO_2_ concentrations, with an average mortality rate of 45% compared to the 69% average mortality found in ambient chambers. See S5 Fig for a comparison of mortality and CO_2_ concentration over time.

### 3.3 LEAF MASS

The effects of the environmental conditions studied in this experiment on lettuce leaf dry biomass are compiled in Figs 8 and 9.

**Figure 8:**
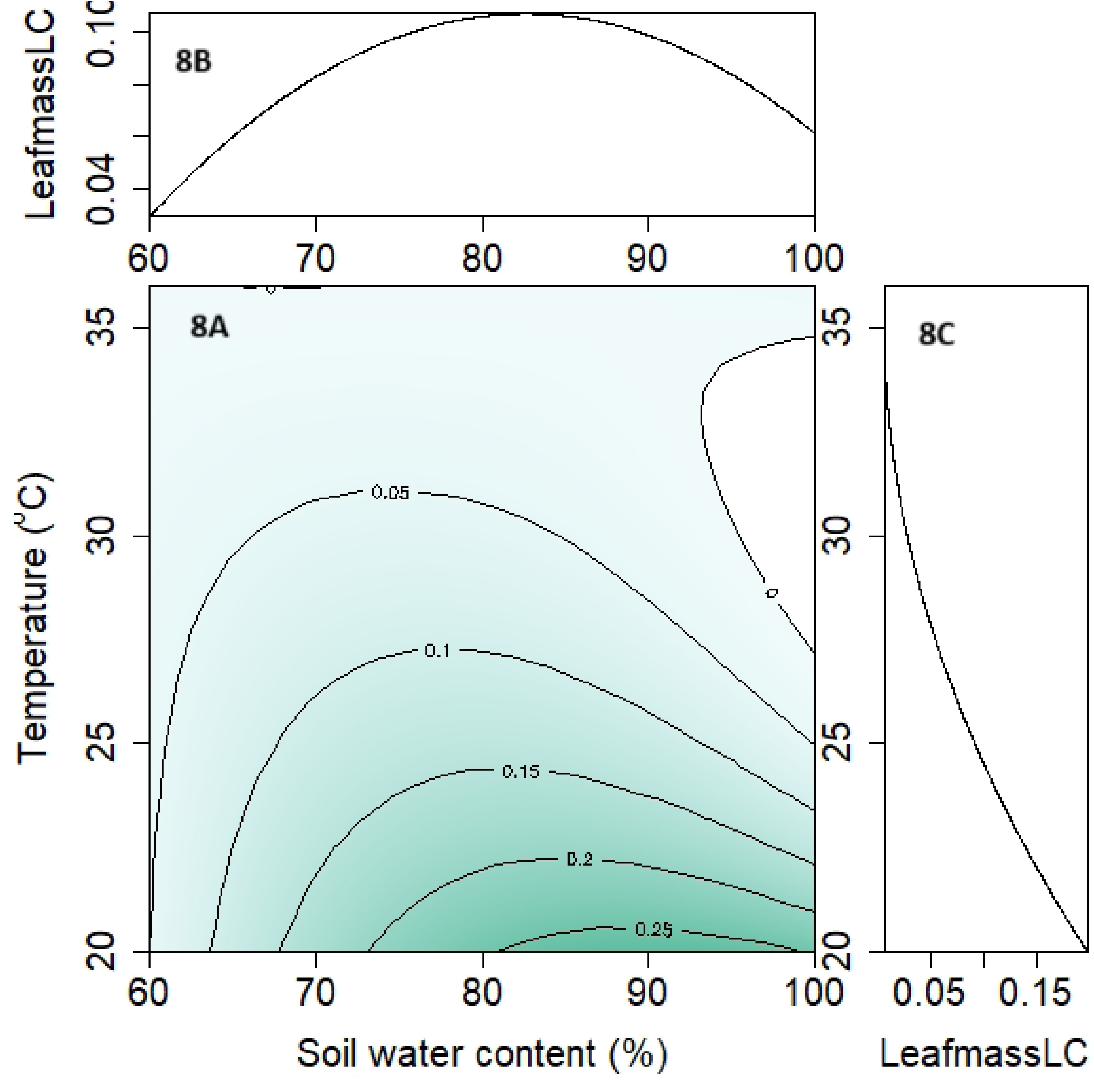
The impacts of temperature and water availability on lettuce dry leaf mass under ambient (LC) and elevated (HC) CO_2_ concentrations. The central figures (8A) show the combined impacts of temperature and water availability on lettuce leaf mass, with darker colors representing higher leaf masses. See contour lines for average dry leaf mass, which is given in grams. The upper subplots (8B) show the relationship between leaf mass and water availability alone and the lower subplots (8C) show the relationship between leaf mass and temperature alone.

**Figure 9:**
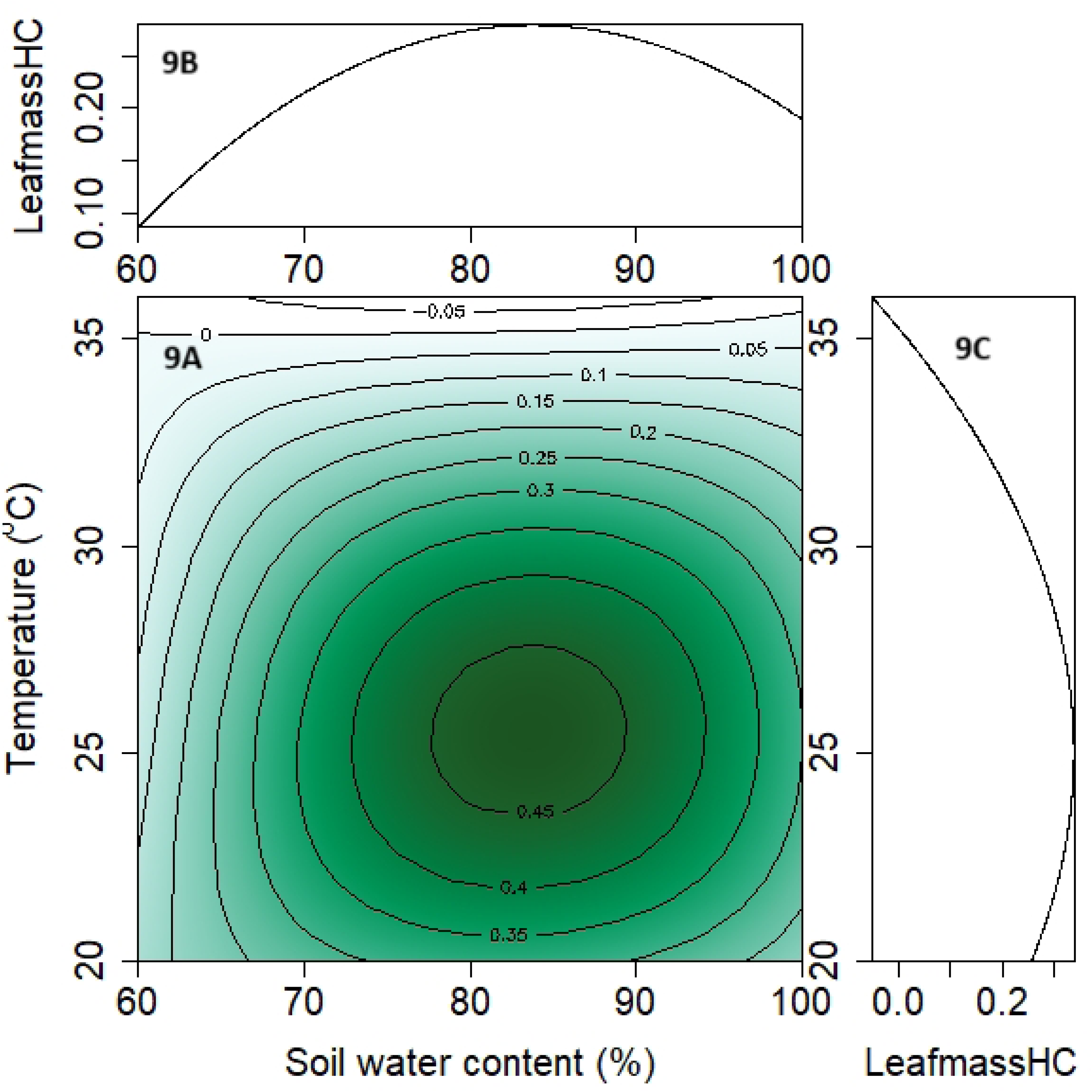
The impacts of temperature and water availability on lettuce dry leaf mass under ambient (LC) and elevated (HC) CO_2_ concentrations. The central figures (9A) show the combined impacts of temperature and water availability on lettuce leaf mass, with darker colors representing higher leaf masses. See contour lines for average dry leaf mass, which is given in grams. The upper subplots (9B) show the relationship between leaf mass and water availability alone and the lower subplots (9C) show the relationship between leaf mass and temperature alone.

In the ambient CO_2_ chambers, leaf mass was highest at low temperatures and medium-high water availability. Generally speaking, temperature had a greater effect on leaf mass than water availability in the ambient chambers. In the elevated CO_2_ chambers, leaf mass was highest at medium temperature and water availability. In the elevated chambers, both temperature and water availability had large impacts on leaf mass. Between the two groups of chambers, leaf mass was generally higher under elevated than ambient CO_2_ concentrations.

#### 3.3.1 RELATIONSHIP WITH TEMPERATURE

In ambient chambers, the average dry leaf mass of surviving plants decreased with increasing temperature. The highest average leaf mass of plants grown in ambient chambers was 0.346 g in the 20 °C chamber, a value that consistently decreased with increasing temperature until reaching 0 g in the 36 °C chamber. When including dead plants - which are considered to have a leaf mass of zero - in the data, leaf mass still declined with increasing temperature, but with less consistency. Most notably, there is a large dip in average leaf mass at 24 °C due to high mortality, even though the average mass of living 24 °C plants was the second highest of all plants grown in ambient chambers.

In elevated chambers, the average dry leaf mass of surviving plants was highest in the 24 °C chamber (0.510 g) and decreased with increasing temperature until again reaching 0 g at 36 °C. This trend was generally mirrored when including dead plants in the calculations, although there is a dip in the data at 32 °C due to high mortality. When comparing chambers with elevated CO_2_ concentrations with ambient chambers set at the same temperature, almost all elevated chambers have larger average leaf masses, regardless of whether dead plants are included or not. See Table 3 and S6 and S7 Figs for more information on the relationship between temperature, average dry leaf mass, and CO_2_ concentration.

**Table 3:** Average leaf masses of elevated and ambient chambers set at different temperatures. When included in the average, dead plants were given a leaf mass of 0 g.

|  | Ambient CO<sub>2</sub> |  | Elevated CO<sub>2</sub> |  |
| --- | --- | --- | --- | --- |
| Temperature (°C) | Average dry leaf mass (g) | Average dry leaf mass (g), including dead plants | Average dry leaf mass (g) | Average dry leaf mass (g), including dead plants |
| 20 °C | 0.346 | 0.228 | 0.312 | 0.205 |
| 24 °C | 0.221 | 0.025 | 0.510 | 0.437 |
| 28 °C | 0.196 | 0.118 | 0.401 | 0.309 |
| 32 °C | 0.076 | 0.009 | 0.304 | 0.078 |
| 36 °C | 0.000 | 0.000 | 0.000 | 0.000 |

#### 3.3.2 RELATIONSHIP WITH SOIL WATER CONTENT

In chambers with ambient CO_2_ concentrations, dry leaf biomass generally increased with increasing soil water content, with the highest average biomass (0.354 g) occurring at 100% field capacity and the lowest (0.136 g) occurring at 60% field capacity. However, when mortality was taken into consideration by including zero values for dead plants, the highest average biomass (0.146 g) occurred at 80% field capacity. Lettuce grown in chambers with elevated CO_2_ followed a similar pattern, with the highest dry biomass occurring at 80% field capacity regardless of whether mortality is considered in the calculation or not. In both ambient and elevated chambers and whether or not mortality was included, plants grown at 60% field capacity always had a lower average dry biomass than plants grown at 100% field capacity. See Table 4 and S8 and S9 Figs for information on the relationship between soil water availability, average dry leaf mass, and CO_2_ concentration.

**Table 4:** Average dry leaf masses of elevated and ambient chambers set at different soil water contents. When included in the average, dead plants are given a leaf mass of 0 g.

| Soil water content (%) | Ambient CO<sub>2</sub> |  | Elevated CO<sub>2</sub> |  |
| --- | --- | --- | --- | --- |
|  | Average dry leaf mass (g) | Average dry leaf mass (g), including dead plants | Average dry leaf mass (g) | Average dry leaf mass (g), including dead plants |
| 100 | 0.354 | 0.061 | 0.411 | 0.188 |
| 90 | 0.278 | 0.087 | 0.512 | 0.249 |
| 80 | 0.341 | 0.146 | 0.528 | 0.332 |
| 70 | 0.165 | 0.042 | 0.318 | 0.154 |
| 60 | 0.136 | 0.043 | 0.220 | 0.107 |

#### 3.3.3 RELATIONSHIP WITH CO_2_ CONCENTRATION

Under elevated CO_2_ concentrations, lettuce plants that survived until the end of the experiment had an average dry leaf biomass that was approximately 59% greater than that of plants grown under ambient concentrations. This difference became more prominent when mortality was considered, as the average dry leaf biomass of elevated CO_2_ plants was over 171% greater than that of ambient plants when dead plants were included as zero values. See Table 5 for information on the relationship between average dry leaf mass and CO_2_ concentration.

**Table 5.** Total and average leaf masses of elevated and ambient chambers. When included in the average, dead plants are given a leaf mass of 0 g. Total Dry Leaf Mass refers to the sum of the dry leaf mass of all surviving plants grown under ambient or elevated CO_2_ concentrations.

|  | Total Dry Leaf Mass (g) | Average Dry Leaf Mass (g) | Average Dry Leaf Mass (g), including dead plants |
| --- | --- | --- | --- |
| Ambient CO<sub>2</sub> | 13.278 | 0.255 | 0.076 |
| Elevated CO<sub>2</sub> | 36.032 | 0.405 | 0.206 |

### 3.4 NITROGEN CONTENT

The effects of the environmental conditions studied in this experiment on lettuce leaf nitrogen content are compiled in Figs 10 & 11.

**Fig 10.**
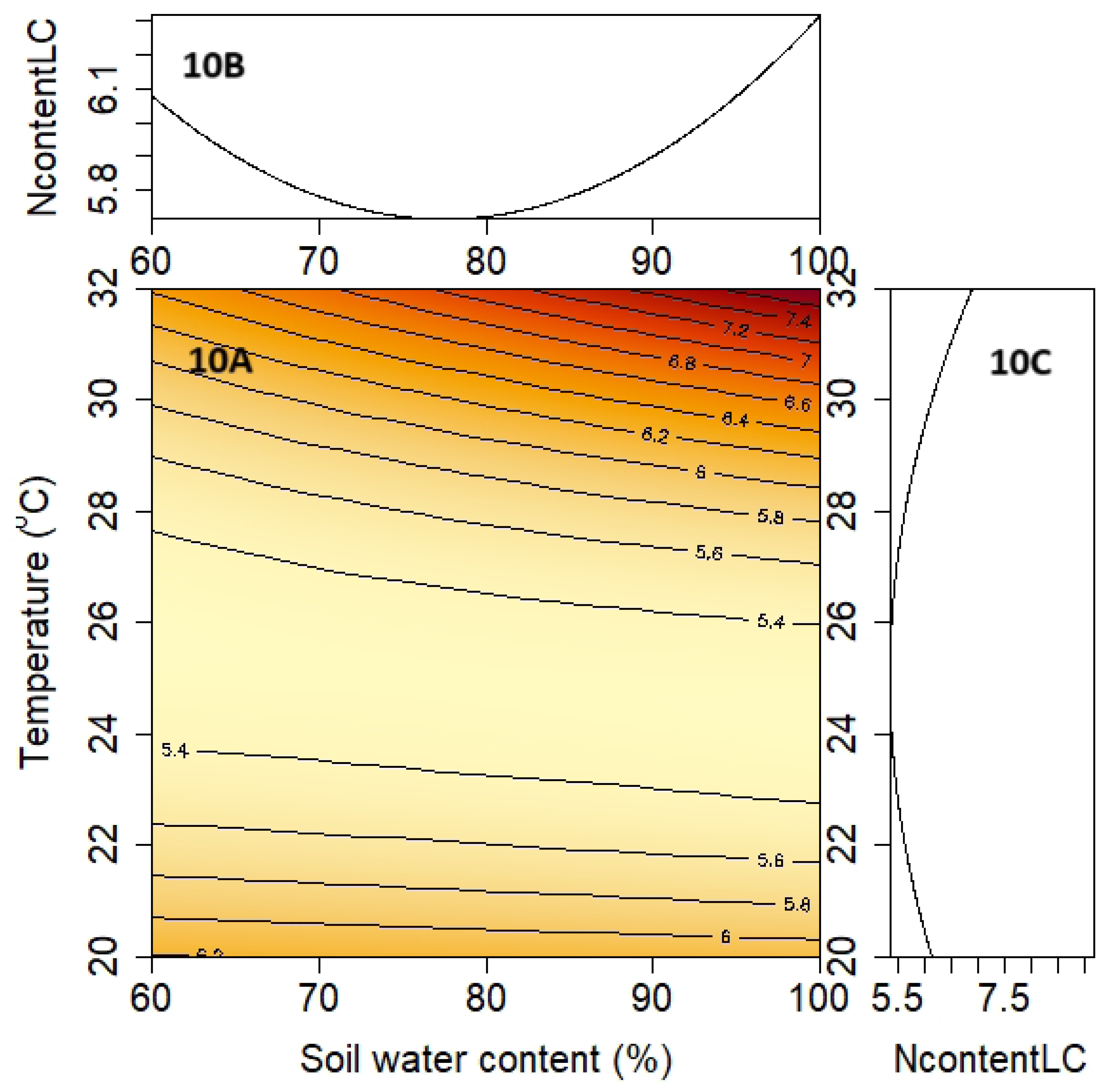
The impacts of temperature and water availability on lettuce nitrogen content under ambient (LC) and elevated (HC) CO_2_ concentrations. The central figures (10A) show the combined impacts of temperature and water availability on lettuce percent nitrogen content, with darker colors representing higher nitrogen content. See contour lines for average lettuce nitrogen content, which is given in percentage of total dry biomass. The upper subplots (10B) show the relationship between nitrogen content and water availability alone and the lower subplots (10C) show the relationship between leaf mass and temperature alone. These figures do not include nitrogen data from the 36 °C chambers, as no living plants remained in said chambers by the end of the experiment.

**Fig 11.**
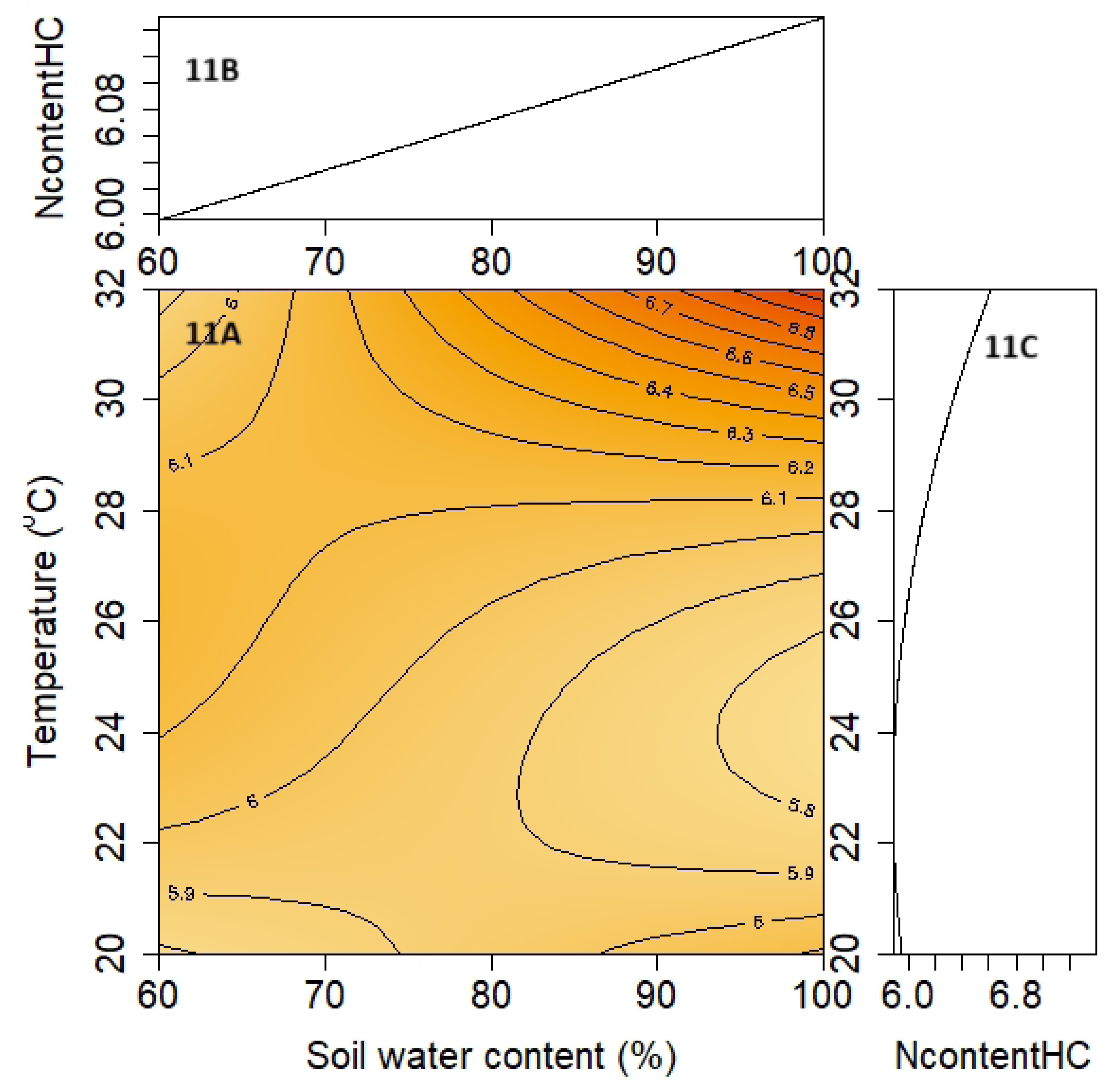
The impacts of temperature and water availability on lettuce nitrogen content under ambient (LC) and elevated (HC) CO_2_ concentrations. The central figures (11A) show the combined impacts of temperature and water availability on lettuce percent nitrogen content, with darker colors representing higher nitrogen content. See contour lines for average lettuce nitrogen content, which is given in percentage of total dry biomass. The upper subplots (11B) show the relationship between nitrogen content and water availability alone and the lower subplots (11C) show the relationship between leaf mass and temperature alone. These figures do not include nitrogen data from the 36 °C chambers, as no living plants remained in said chambers by the end of the experiment.

In Fig 22, the highest leaf nitrogen content was found at high temperature and water availability. This decreased under medium temperatures before slightly increasing again at the lowest temperature. In general, temperature had a more important effect on nitrogen content than water availability. In Fig 23, the highest nitrogen levels were again found at high temperatures and water availability. Temperature again generally had a more important effect than water availability. Between the two figures, there is little average difference in nitrogen content between ambient and elevated CO_2_ concentrations, with elevated chambers leading to slightly higher nitrogen content.

#### 3.4.1 RELATIONSHIP WITH TEMPERATURE

In regard to temperature, the highest average percent nitrogen found in dry leaf samples occurred in the 32 °C chambers for both ambient and elevated CO_2_. These values - 6.92% and 6.76% respectively - were also the highest average nitrogen content values found in any of the chambers. In the ambient chambers, N content decreased with decreasing temperature until increasing again at 20 °C. The lowest average N content of all chambers (5.44%) was found at 20 °C and ambient CO_2_. The elevated chambers expressed a less definite relationship between temperature and N content, with a decrease in N content when temperatures decreased from 32 °C to 28 °C but few changes in N content between the other three chambers. See Table 6 and S10 Fig for information on the relationship between temperature, CO_2_ concentration, and nitrogen content.

**Table 6:** Average nitrogen content of dry lettuce leaf biomass grown under different temperatures and under ambient and elevated CO_2_.

| Temperature (°C) | Average Nitrogen Content Under Ambient CO<sub>2</sub> (%) | Average Nitrogen Content Under Elevated CO<sub>2</sub> (%) |
| --- | --- | --- |
| 20 | 6.14 | 5.89 |
| 24 | 5.44 | 6.10 |
| 28 | 5.63 | 5.92 |
| 32 | 6.92 | 6.76 |

#### 3.4.2 RELATIONSHIP WITH WATER AVAILABILITY

Within the elevated chambers, there was a small increase in N content with increasing soil water content. The highest average N content found in elevated chambers (6.158%) was at 100% field capacity and the lowest (5.992%) was found at 70%. There was less of a trend within ambient chambers, as the highest value (6.27%) was found at 90% field capacity and the smallest (5.53%) was found at 80%. See Table 7 and S11 Fig for more information on the relationship between water availability, CO_2_ concentration, and nitrogen content.

**Table 7:** Average nitrogen content of dry lettuce leaf biomass grown under different levels of water availability and under ambient and elevated CO_2_.

| Soil Water Content (%) | Average N Content (%) Under Ambient CO<sub>2</sub> | Average N Content (%) Under Elevated CO<sub>2</sub> |
| --- | --- | --- |
| 100 | 6.098 | 6.158 |
| 90 | 6.286 | 6.071 |
| 80 | 5.531 | 6.123 |
| 70 | 5.670 | 5.992 |
| 60 | 6.196 | 6.008 |

#### 3.4.3 RELATIONSHIP WITH CO_2_

CO_2_ concentration had little effect on lettuce leaf nitrogen content. On average, plants grown under ambient CO_2_ concentration had a dry leaf biomass that was 5.93% nitrogen and plants grown under elevated concentrations had an average dry leaf biomass that was 6.08% nitrogen.

## 4.0 DISCUSSION

### 4.1 IMPLICATIONS OF CLIMATE CHANGE IMPACTS ON LETTUCE GROWTH AND MORTALITY

Because of the wide variety of environments lettuce is grown under and the differing ways that these environments could be impacted by climate change, it is difficult to make any concrete, overarching predictions as to how climate change could affect global lettuce harvests. With this being said, our research generally suggests that climate change has the potential to improve lettuce growth and survival due to the beneficial effects of elevated CO_2_ concentrations. However, this improvement could be negated by the deleterious effects of high temperatures, drought, and/or flooding. These tentative conclusions are largely supported by similar studies, which are discussed below.

#### 4.1.1 CO_2_ IMPACTS

As predicted by our hypothesis, elevated CO_2_ concentrations led to larger plant biomass and improved plant survival across all treatments. This increase in productivity is likely due primarily to the photosynthesis-promoting effects of increased CO_2_ availability [45–46]. Our data also suggests that increased CO_2_ concentrations improve lettuce thermal tolerance. As shown in Figs 16 and 17, while lettuce biomass increased with decreasing temperature under ambient CO_2_ concentration, under elevated concentrations biomass was greatest at approximately 25 °C. The relationship between biomass and water availability, on the other hand, is roughly the same under both ambient and elevated CO_2_ concentrations. The similar relationship between water availability and biomass in both ambient and elevated treatments suggests that improved water use efficiency due to elevated CO_2_ concentrations does not have a significant effect on lettuce grown under the levels of water availability used in this study.

Comparing our research to the literature shows that the relationship we found between CO_2_ availability and biomass yield is supported by the findings of [27, 45–47], all of whom found that elevated CO_2_ concentrations increased lettuce biomass production. [47] also found that artificially elevating CO_2_ concentrations led to an increase in the optimal temperature for lettuce cultivar biomass growth from 25 °C to 30 °C. However, when comparing the growth rates of lettuce grown under a wide range of CO_2_ concentrations, [45] found that increasing CO_2_ levels gave diminishing returns in biomass increases once concentrations surpassed 800 ppm.

#### 4.1.2 TEMPERATURE IMPACTS

Within both the ambient CO_2_ group and the elevated CO_2_ group, our data suggests that the success of Mānoa lettuce declines rapidly at high temperatures and that the cultivar cannot survive at temperatures exceeding 36 °C. This supports our initial hypothesis, which predicted that lettuce growth would be negatively impacted by increasing temperatures. This could be due to a number of factors, including damaged cellular apparatuses, impaired enzyme function, and the accumulation of reactive oxygen species [8–10]. Biomass and survival both increased at lower temperatures, with the optimum temperature for lettuce grown under elevated CO_2_ falling around 25 °C and the optimum temperature under ambient CO_2_ falling around 20 °C or lower. This is notable, as the Mānoa lettuce cultivar is ordinarily known to grow poorly at temperatures above 20 °C [48].

The relationship between temperature and lettuce growth has been explored by a number of studies, but there is limited consensus as to the nature of this relationship. Studies by [49–50] have found that lettuce biomass increased as temperature increased from 20 °C to 24 °C and 25 °C to 33 °C, respectively. However, studies by [51–52] found that biomass decreased as temperatures increased from 20 °C to 30 °C and 14 °C to 21°C, respectively. This discrepancy could be due to different lettuce cultivars having differing optimal temperatures, as demonstrated by [53–55].

#### 4.1.3 WATER AVAILABILITY IMPACTS

Although temperature had a greater impact on plant survival and yield, water availability also played an important role in lettuce growth and survival. Our findings from both elevated and ambient treatments suggest that the optimum level of substrate water content for Mānoa lettuce grown in coarse vermiculite is approximately 80% of the experimentally determined field capacity of the media. This differs from our hypothesis, which theorized that lettuce growth would be greatest at the highest level of water availability. The decrease in lettuce productivity at lower water availability could be due to lower rates of water flow to new growth [14, 17] or closed stomata reducing the amount of CO_2_ available for photosynthesis [14–16]. At higher water availability, it is possible that waterlogging could lead to root oxygen deprivation, which can have detrimental effects on the uptake of water and nutrients [56–57].

Other studies on the effect of water availability on lettuce have also found that both drought and waterlogging can have negative impacts on lettuce biomass production. When growing lettuce plants under both waterlogging and drought stress, [58] found that waterlogging had a larger negative impact on plant growth than drought. Both [59–60] compared the effects of waterlogging and water restriction on different lettuce cultivars and found that, although both water extremes reduced plant growth, the degree of this impact depended on the cultivar being grown. Additionally, the soil microbial content of a growing media can modify the effects of both drought stress [61–62] and waterlogging [61, 63–64] on plant growth and survival. Thus, the lack of a robust microbial community in the growth medium we used could have further influenced the response of the experimental plants to water availability.

### 4.2 IMPACTS OF CLIMATE CHANGE FACTORS ON NITROGEN CONTENT

Based on our findings, it appears that lettuce percent nitrogen content may slightly increase when both temperature and water availability are at high levels. The data suggests that this increase may be less prominent under elevated CO_2_ concentrations, which could potentially lead to reduced nutritional quality in lettuce under climate change conditions. However, under lower temperature and water availability conditions, the nitrogen contents of lettuce plants grown under ambient and elevated CO_2_ concentration were more similar. These findings partially support our hypothesis that lettuce nitrogen content would decrease under elevated CO_2_ concentrations, but only at high temperatures and with abundant water availability. With this being said, the small sample size of high temperature lettuce that was available for testing reduces the reliability of this prediction.

Our findings regarding the impacts of climate change on lettuce nitrogen content are partially supported by the literature, although few studies have examined the impacts of climate change on lettuce nitrogen content specifically. One study by [27] examined lettuce and spinach plants grown under elevated CO_2_ concentrations and found that increased concentrations typically led to decreased concentrations of nitrogen, potassium, and phosphorus in the leaves of both species. However, a review of 54 studies by [30] found that, although the protein content of almost all crops decreased when grown under elevated CO_2_, leafy vegetables did not experience a significant decrease in protein content. With this being said, the same study also found that increased CO_2_ concentrations led to a decrease in nitrate in leafy vegetables of 18.8%, which could contribute to a decrease in total nitrogen content [30]. In regards to temperature, some studies have concluded that lettuce protein content increases with warming conditions [49] while others have reported the opposite trend [65]. It is also possible that the higher levels of nitrogen under high temperatures and lower water availability could be due to the smaller size of surviving lettuce plants under these conditions, as the reduced amount of nonstructural carbohydrates in smaller plants could lead to a larger quantity of nitrogen per unit of biomass [31, 33–34].

### 4.3 LIMITATIONS

Because our experiment took place under laboratory conditions, there are several areas in which our research conditions differed from the conditions lettuce would grow under in a farm setting. For one, the media used to grow our lettuce was vermiculite, a sterile, inorganic media that lacks the microbiome found in most organic soils. Vermiculite also has a high water capacity that allows it to hold a larger amount of moisture than many organic soils. Additionally, because the pots used in the IPS system lack drainage holes, there was no opportunity for excess water or nutrients to escape from each pot. This lack of drainage has the potential to promote the proliferation of harmful microorganisms and increase waterlogging. Microbial growth could also be encouraged by low ventilation, which promoted high humidity levels inside the chambers. Finally, growing indoors eliminates pressures from pests, weeds, or adverse weather conditions, all of which can influence plant growth in outdoor settings.

Other factors that could have contributed to inaccuracy in our data include CO_2_ fluctuations and laboratory malfunctions. As mentioned previously, the ambient CO_2_ concentrations in the lab were not consistent and depended largely on the number of people present in the building. On an ordinary day, this meant that CO_2_ levels in the laboratory ranged from approximately 430 ppm to 520 ppm. When the lab was actively in use for long periods of time, CO_2_ concentrations could reach as high as 600 ppm. As a result, the average CO_2_ concentration in the ambient chambers was well above the April 2023 atmospheric CO_2_ concentration of 423 ppm [66] recorded by the Mauna Loa observatory. Additionally, the 20 °C chambers - which were the most energy-intensive - experienced two power outages each over the course of the experiment. During these outages, the chambers warmed to room temperature (approximately 25 °C at night), lost all light if the outage occurred during the 12 hour “daytime”, and, in the case of the elevated CO_2_ chamber, returned to ambient CO_2_ concentrations for the duration of the outage. Although only two of these events occurred during the experiment and both lasted for less than 6 hours, power loss could have impacted lettuce growth rates.

### 4.4 POSSIBLE FUTURE RESEARCH

There are multiple possible options for future research into the effects of climate change on lettuce and other plants. Because this study only examined the upper ends of the survivable spectrums of temperature and water availability, future research could explore the lower limits as well. Studying the lower limit of water availability under ambient and elevated CO_2_ is especially important, as elevated CO_2_ should theoretically allow lettuce to survive under drier conditions due to improved water use efficiency. For similar reasons, future research could explore the responses of lettuce plants to different nutrient concentrations. Plant nutrient uptake can be influenced by temperature, CO_2_ concentration, and water stress, so it is fully possible that climate change could change the way lettuce interacts with soil nutrients [67–68]. Nutrient uptake can also be influenced by the soil microbiome, another factor that could be incorporated into future studies [61, 63]. Finally, there is a great need for more research into the effects of climate change on other lettuce cultivars as well as other food crops. Climate change impacts are highly species- and cultivar-specific, so the results of one study on one lettuce cultivar are not enough to provide accurate insight into the effects of environmental change on the global food system [22].

## 5.0 CONCLUSION

It is highly likely that climate change will affect the global agricultural system, but the limited information available on the effects of climate change on crop plants can make it difficult to predict how future agricultural landscapes could function. To investigate the impacts of climate change on one particular crop - lettuce - we grew Mānoa lettuce under a wide variety of environmental conditions and observed the way lettuce growth was modified by changing conditions. We found that, although lettuce growth and survival was improved by increasing atmospheric CO_2_ concentrations, this benefit could be reduced or negated by the detrimental effects of high atmospheric temperatures and high or low water availability. Additionally, we found that climate change conditions have the potential to impact the nitrogen content of lettuce and may decrease protein content under high temperatures and waterlogged conditions. This information could be very useful for the lettuce farmers of today, as it suggests that lettuce yield could be improved in the future, but only if the area in which it is grown is not expected to experience significant temperature increases or increases or decreases in precipitation. Moving forward, it is vital that future studies explore the potential impacts of climate change on other plant and animal species. In order to adequately prepare for a future where climate change may radically alter the way organisms grow and function, the scientific community of today must invest time and resources into understanding the potential landscapes of tomorrow.

## Supporting figure and table captions

**S1 Table. The six models used in the regression analysis for this study, including AIC, R^2^, and p-values.**

**S2 Table. P-values for the individual predictor variables for the six models.**

**S1 Fig. Percent mortality over time (days) at different temperatures under ambient CO_2_ conditions.**

**S2 Fig. Percent mortality over time (days) at different temperatures under elevated CO_2_ conditions.**

**S3 Fig. Percent mortality over time (days) for chambers set at different levels of soil water availability (% of maximum field capacity) under ambient CO_2_ concentrations.**

**S4 Fig. Percent mortality over time (days) for chambers set at different levels of soil water availability (% of maximum field capacity) under elevated CO_2_ concentrations.**

**S5 Fig. Average percent mortality over time (days) for lettuce grown under ambient and elevated CO_2_ concentrations.**

**S6 Fig. Average dry lettuce leaf mass (g) versus chamber temperature (°C) for plants grown under ambient and elevated CO_2_ concentrations.** This figure only considers the average masses of plants that survived until the end of the experiment.

**S7 Fig. Average dry lettuce leaf mass (g) versus chamber temperature (°C) for plants grown under ambient and elevated CO_2_ concentrations, incorporating mortality.** This figure considers both the average masses of plants that survived until the end of the experiment as well as the masses of plants that did not survive, the latter of which are considered to have a mass of 0 g.

**S8 Fig. Average dry lettuce leaf mass (g) versus chamber soil water availability (% of field capacity) for plants grown under ambient and elevated CO_2_ concentrations.** This figure only considers the average masses of plants that survived until the end of the experiment.

**S9 Fig. Average dry lettuce leaf mass (g) versus chamber soil water availability (% of field capacity) for plants grown under ambient and elevated CO_2_ concentrations, incorporating mortality.** This figure considers both the average masses of plants that survived until the end of the experiment as well as the masses of plants that did not survive, the latter of which are considered to have a mass of 0 g.

**S10 Fig. Line graph showing average dry lettuce leaf N content (%) versus chamber temperature (°C) for plants grown under ambient and elevated CO_2_ concentrations.** The solid line represents the average nitrogen content of dry leaf samples grown under ambient CO_2_ levels while the dotted line represents the same data for plants grown under elevated CO_2_ levels. This figure does not include nitrogen data from the 36 °C chambers, as no living plants remained in said chambers by the end of the experiment.

**S11 Fig. Line graph showing average dry lettuce leaf N content (%) versus chamber soil water availability (%) for plants grown under ambient and elevated CO_2_ concentrations.** The solid line represents the average nitrogen content of dry leaf samples grown under ambient CO_2_ levels while the dotted line represents the same data for plants grown under elevated CO_2_ levels.

## Notes

### Competing Interest Statement

The authors have declared no competing interest.

## LITERATURE CITED

1. Janowiak M, Connelly WJ, Dante-Wood K, Domke GM, Giardina C, Kayler Z, et al. Considering Forest and Grassland Carbon in Land Management. Washington, D.C.: U.S. Department of Agriculture, Forest Service; 2017. doi: 10.2737%2Fwo-gtr-95

2. Zuazo VHD, Pleguezuelo CRR. Soil-Erosion and Runoff Prevention by Plant Covers: A Review. In: Lichtfouse E, Navarrete M, Debaeke P, Véronique S, Alberola C, editors. Sustainable Agriculture. Dordrecht: Springer; 2009. p. 785–811. doi: 10.1007/978-90-481-2666-8_48

3. Ernst C, Gullick R, Nixon K. Conserving Forests to Protect Water. Opflow. 2004;30(5):1–7. doi: 10.1002/j.1551-8701.2004.tb01752.x

4. Perkins KS, Nimmo JR, Medeiros AC, Szutu DJ, von Allmen E. Assessing effects of native forest restoration on soil moisture dynamics and potential aquifer recharge, Auwahi, Maui. Ecohydrology. 2004;7(5):1437–1451. doi: 10.1002/eco.1469

5. Food and Agriculture Organization of the United Nations (FAO). FAOSTAT [Internet]. Rome: Food and Agriculture Organization of the United Nations (IT). [cited 2024 Jun 11]. Available from: https://www.fao.org/faostat/en/#data

6. United States Census Bureau (USCB). Highlights of 2023 Characteristics of New Housing. 2024 Jun 3 [cited 2024 Jun 17] In: USCB website [Internet]. Suidland (MD): UCSB - . [about 2 screens] Available from: https://www.census.gov/construction/chars/highlights.html

7. Raza A, Razzaq A, Mehmood SS, Zou X, Zhang X, Lv Y, et al. Impact of Climate Change on Crops Adaptation and Strategies to Tackle Its Outcome: A Review. Plants. 2019;8(2):34. doi: 10.3390/plants8020034

8. Bita CE, Gerats T. Plant tolerance to high temperature in a changing environment: Scientific fundamentals and production of heat stress-tolerant crops. Front Plant Sci. 2013;4(273). doi: 10.3389/fpls.2013.00273

9. Feller U, Vaseva II. Extreme climatic events: impacts of drought and high temperature on physiological processes in agronomically important plants. Front Environ Sci. 2014;2. doi: 10.3389/fenvs.2014.00039

10. Hasanuzzaman M, Nahar K, Alam M, Roychowdhury R, Fujita M. Physiological, Biochemical, and Molecular Mechanisms of Heat Stress Tolerance in Plants. Int J Mol Sci. 2013;14(5):9643–84. doi: 10.3390/ijms14059643

11. Song Y, Chen Q, Ci D, Shao X, Zhang D. Effects of high temperature on photosynthesis and related gene expression in poplar. BMC Plant Biol. 2014:14:111. doi:10.1186/1471-2229-14-111

12. Nafees M, Fahad S, Shah AN, Bukhari MA, Maryam, Ahmed I, et al. Reactive Oxygen Species Signaling in Plants. In: Hasanuzzaman M, Hakeem K, Nahar K, Alharby H, editors. Plant Abiotic Stress Tolerance. Cham: Springer; 2019. p. 259–272. doi: 10.1007/978-3-030-06118-0_11

13. Lamaoui M, Jemo M, Datla R, Bekkaoui F. Heat and Drought Stresses in Crops and Approaches for Their Mitigation. Front Chem. 2018;2. doi: 10.3389/fchem.2018.00026

14. Kaur G, Asthir B. Molecular responses to drought stress in plants. Biol Plant. 2017;61:201–209. doi: 10.1007/s10535-016-0700-9

15. Lawlor DW. Limitation to Photosynthesis in Water-stressed Leaves: Stomata vs. Metabolism and the Role of ATP. Ann Bot. 2002;89(7):871–885. doi: 10.1093/aob/mcf110

16. Osakabe Y, Osakabe K, Shinozaki K, Tran L. Response of plants to water stress. Front Plant Sci. 2014;5:86. doi: 10.3389/fpls.2014.00086

17. Seleiman MF, Al-Suhaibani N, Ali N, Akmal M, Alotaibi M, Refay Y, et al. Drought Stress Impacts on Plants and Different Approaches to Alleviate Its Adverse Effects. Plants. 2021;10(2):259. doi: 10.3390/plants10020259

18. Cernusak LA, Haverd V, Brende O, Thiec DL, Guehl J, Cuntz M. Robust Response of Terrestrial Plants to Rising CO_2_. Trends Plant Sci. 2019;24(7), 578–586. doi: 10.1016/j.tplants.2019.04.003

19. Amthor JS. Terrestrial higher-plant response to increasing atmospheric [CO_2_] in relation to the global carbon cycle. Glob Chang Biol. 1995;1(4):243–274. doi: 10.1111/j.1365-2486.1995.tb00025.x

20. Ackerly DD, Bazzaz FA. Plant growth and reproduction along CO_2_ gradients: non-linear responses and implications for community change. Glob Chang Biol. 1995;1(3):199–207. doi: 10.1111/j.1365-2486.1995.tb00021.x

21. Díaz S, Grime J, Harris J, McPherson E. Evidence of a feedback mechanism limiting plant response to elevated carbon dioxide. Nature. 1993;364:616–617. doi: 10.1038/364616a0

22. Parmesan C, Hanley ME. Plants and climate change: complexities and surprises. Ann Bot. 2015;116(6):849–864. doi: 10.1093/aob/mcv169

23. Urban O. Physiological Impacts of Elevated CO_2_ Concentration Ranging from Molecular to Whole Plant Responses. Photosynthetica. 2003;41:9–20. doi: 10.1023/A:1025891825050

24. Damour G, Simonneau T, Cochard H, Urban L. An overview of models of stomatal conductance at the leaf level. Plant Cell Environ. 2010;33(9):1419–1438. doi: 10.1111/j.1365-3040.2010.02181.x

25. Xu Z, Shimizu H, Yagasaki Y, Ito S, Zheng Y, Zhou G. Interactive Effects of Elevated CO_2_, Drought, and Warming on Plants. J Plant Growth Regul. 2013;32:692–707. doi: 10.1007/s00344-013-9337-5

26. Urban J, Ingwers M, McGuire MA, Teskey RO. Stomatal conductance increases with rising temperature. Plant Signal Behav. 2017;12(8):e1356534. doi: 10.1080/15592324.2017.1356534

27. Giri A, Armstrong B, Rajashekar CB. Elevated Carbon Dioxide Level Suppresses Nutritional Quality of Lettuce and Spinach. Am J Plant Sci. 2016;7(1):246–258. doi: 10.4236/ajps.2016.71024

28. DaMatta FM, Grandis A, Arenque BC, Buckeridge MS. Impacts of climate changes on crop physiology and food quality. Food Res Int. 2010;43(7):1814–1823. doi: 10.1016/j.foodres.2009.11.001.

29. Taub DR, Miller B, Allen H. Effects of elevated CO2 on the protein concentration of food crops: a meta-analysis. Glob Chang Biol. 2007;14(3):565–575. doi: 10.1111/j.1365-2486.2007.01511.x

30. Dong J, Gruda N, Lam SK, Li X, Duan Z. Effects of Elevated CO_2_ on Nutritional Quality of Vegetables: A Review. Front Plant Sci. 2018;9:924. doi: 10.3389/fpls.2018.00924

31. Idso SB, Idso KE. Effects of atmospheric CO_2_ enrichment on plant constituents related to animal and human health. Environ Exp Bot. 2001;45(2):179–199. doi: 10.1016/S0098-8472(00)00091-5

32. Kuehny JS, Peet MM, Nelson PV, Willits D. Nutrient Dilution by Starch in CO_2_-enriched Chrysanthemum. J Exp Bot. 1991;42(6):711–716. doi: 10.1093/jxb/42.6.711

33. Taub DR, Wang X. Why are Nitrogen Concentrations in Plant Tissues Lower under Elevated CO_2_? A Critical Examination of the Hypotheses. J Integr Plant Biol. 2008;50(11):1365–1374. doi: 10.1111/j.1744-7909.2008.00754.x

34. Peet MM, Wolfe DW. Crop Ecosystems Responses to Climatic Change: Vegetable Crops. In: Reddy KR, Hodges HF, editors. Climate Change and Global Crop Productivity Wallingford: CABI Publishing; 2000. p. 213–244. doi: 10.1079/9780851994390.0213

35. Stitt M, Krapp A. The interaction between elevated carbon dioxide and nitrogen nutrition: the physiological and molecular background. Plant Cell Environ. 2002;22(6):583–621. doi: 10.1046/j.1365-3040.1999.00386.x

36. Conroy J, Hocking P. Nitrogen nutrition of C_3_ plants at elevated atmospheric CO_2_ concentrations. Physiol Plant. 1993;89(3):570–576. doi: 10.1111/j.1399-3054.1993.tb05215.x

37. Shatilov M, Razin A, Ivanova M. Analysis of the world lettuce market. IOP Conf Ser Earth Environ Sci. 2019;395:012053. doi: 10.1088/1755-1315/395/1/012053

38. Holmes SC, Wells DE, Pickens JM, Kemble JM. Selection of Heat Tolerant Lettuce (*Lactuca sativa* L.) Cultivars Grown in Deep Water Culture and Their Marketability. Horticulturae. 2019;5(3):50. doi: 10.3390/horticulturae5030050

39. Jiménez-Arias, D., García-Machado, F. J., Morales-Sierra, S., Luis, J. C., Suarez, E., Hernández, et al. Lettuce plants treated with L-pyroglutamic acid increase yield under water deficit stress. Environ Exp Bot. 2019;158:215–222. doi: 10.1016/j.envexpbot.2018.10.034

40. Webster KM. Reconciling conflicting results regarding climate change effects on plants: A case study with wheat. M.Sc. Thesis. Mānoa (HI): University of Hawai‘i at Mānoa; 2021. Available from: http://hdl.handle.net/10125/75936

41. McDowell K, Zhong Y, Webster K, Gonzalez HJ, Trimble AZ, Mora C. Comprehensive temperature controller with internet connectivity for plant growth experiments. HardwareX. 2021;10:e00238. doi: 10.1016/j.ohx.2021.e00238

42. Cassel DK, Nielsen DR. Field capacity and available water capacity. In: Klute, A. (eds). Methods of soil analysis: Part 1 Physical and mineralogical methods. Madison: American Society of Agronomy; 1986. p. 901–926. doi: 10.2136/sssabookser5.1.2ed.c36

43. Takara G, Trimble AZ, Arata R, Brown S, Gonzalez HJ, Mora C. An inexpensive robotic gantry to screen and control soil moisture for plant experiments. HardwareX. 2021;9:e00174. doi: 10.1016/j.ohx.2021.e00174

44. Intergovernmental Panel on Climate Change (IPCC). Technical Summary. In: Climate Change 2021 – The Physical Science Basis: Working Group I Contribution to the Sixth Assessment Report of the Intergovernmental Panel on Climate Change. Cambridge (UK): Cambridge University Press; 2021. p. 35–144. doi: 10.1017/9781009157896.002

45. Holley J, Mattson N, Ashenafi E, Nyman M. The Impact of CO_2_ Enrichment on Biomass, Carotenoids, Xanthophyll, and Mineral Content of Lettuce (Lactuca sativa L.). Horticulturae. 2021;8(9):820. doi: 10.3390/horticulturae8090820

46. Pérez-López U, Miranda-Apodaca J, Lacuesta M, Mena-Petite A, Muñoz-Rueda A. Growth and nutritional quality improvement in two differently pigmented lettuce cultivars grown under elevated CO_2_ and/or salinity. Scientia Horticulturae. 2015;195:56–66. doi: 10.1016/j.scienta.2015.08.034

47. Frantz JM, Ritchie G, Cometti NN, Robinson J, Bugbee B. Exploring the Limits of Crop Productivity: Beyond the Limits of Tipburn in Lettuce. J Am Soc Hortic Sci. 2004;129(3):331–338. doi: 10.21273/JASHS.129.3.0331

48. Valenzuela HR, Kratky B, Cho J. Lettuce production guidelines for Hawaii. Honolulu (HI): University of Hawaii; 1996.

49. Chen Z, Jahan MS, Mao P, Wang M, Liu X, Guo S. Functional growth, photosynthesis and nutritional property analyses of lettuce grown under different temperature and light intensity. The Journal of Horticultural Science and Biotechnology, 2020;96(1):53–61. doi: 10.1080/14620316.2020.1807416

50. Sublett WL, Barickman TC, Sams CE. Effects of Elevated Temperature and Potassium on Biomass and Quality of Dark Red ‘Lollo Rosso’ Lettuce. Horticulturae. 2018;4(2):11. doi: 10.3390/horticulturae4020011

51. Iqbal Z, Munir M, Sattar MN. Morphological, Biochemical, and Physiological Response of Butterhead Lettuce to Photo-Thermal Environments. Horticulturae. 2022;8:515. doi: 10.3390/horticulturae8060515

52. Wheeler TR, Hadley P, Morison JIL, Ellis RH. Effects of temperature on the growth of lettuce (Lactuca sativa L.) and the implications for assessing the impacts of potential climate change. Euro J Agron. 1993;2(4):305–311. doi: 10.1016/S1161-0301(14)80178-0

53. Smeets L. Analysis of the differences in growth between five lettuce cultivars marking the development in lettuce breeding for winter production. Euphytica. 1977;26:655–659. doi: 10.1007/BF00021690

54. Choi KY, Paek KY, Lee YB. Effect of Air Temperature on Tipburn Incidence of Butterhead and Leaf Lettuce in a Plant Factory. In: Kubota C, Chun C, editors. Transplant Production in the 21st Century. Dordrecht: Springer; 2000. p . 166–171. doi: 10.1007/978-94-015-9371-7_27

55. Thakulla D, Dunn B, Hu B, Goad C, Maness N. Nutrient Solution Temperature Affects Growth and °Brix Parameters of Seventeen Lettuce Cultivars Grown in an NFT Hydroponic System. Horticulturae. 2021;7(9):321. doi: 10.3390/horticulturae7090321

56. Sairam RK, Kumutha D, Ezhilmathi K, Deshmukh PS, Srivastava GC. Physiology and biochemistry of waterlogging tolerance in plants. Biol Plant. 2008;52:401–412. doi: 10.1007/s10535-008-0084-6

57. Manik SMN, Pengilley G, Dean G, Field B, Shabala S, Zhou M. Soil and Crop Management Practices to Minimize the Impact of Waterlogging on Crop Productivity. Front Plant Sci. 2019;10:140. doi: 10.3389/fpls.2019.00140

58. Cabillo CM. Biomass Production of Lettuce (*Lactuca sativa* L.) Under Water Stress. Int J Sci Eng Res. 2019;10(12): 928–934.

59. Eichholz I, Förster N, Ulrichs C, Schreiner M, Huyskens-Keil S. Survey of bioactive metabolites in selected cultivars and varieties of Lactuca sativa L. under water stress. J Appl Bot Food Qual. 2014;87:265–273. doi: 10.5073/JABFQ.2014.087.037

60. Trang NTD, Schierup H, Brix H. Leaf vegetables for use in integrated hydroponics and aquaculture systems: Effects of root flooding on growth, mineral composition and nutrient uptake. Afr J Biotechnol. 2010;9(27):4186–4196.

61. Ruiz-Lozano JM, Aroca R, Zamarreño AM, Molina S, Andreo-Jiméne, B, Porcel R, et al. Arbuscular mycorrhizal symbiosis induces strigolactone biosynthesis under drought and improves drought tolerance in lettuce and tomato. Plant Cell Environ. 2016;39(2):441–452. doi: 10.1111/pce.12631

62. Durán P, Acuña JJ, Armada E, López-Castillo OM, Cornejo P, Mora ML, et al. Inoculation with selenobacteria and arbuscular mycorrhizal fungi to enhance selenium content in lettuce plants and improve tolerance against drought stress. J Soil Sci Plant Nutr. 2016;16(1):201–225. doi:10.4067/S0718-95162016005000017

63. Irfan M, Hayat S, Hayat Q, Afroz S, Ahmad A. Physiological and biochemical changes in plants under waterlogging. Protoplasma. 2010;241(1-4):3–17. doi: 10.1007/s00709-009-0098-8

64. Laanbroek HJ. Bacterial cycling of minerals that affect plant growth in waterlogged soils: a review. Aquat Bot. 1990;38(1):109–125. doi: 10.1016/0304-3770(90)90101-P

65. Ouyang Z, Tian J, Yan X, Shen H. Effects of different concentrations of dissolved oxygen or temperatures on the growth, photosynthesis, yield and quality of lettuce. Agric Water Manag. 2020;228:105896. doi: 10.1016/j.agwat.2019.105896

66. National Oceanic and Atmospheric Administration (NOAA) Global Monitoring Laboratory. Trends in CO_2_, CH_4_, N_2_O, SF_6_. 2024 June 5 [cited 21 June 2024]. In: NOAA Website [Internet]. Silver Spring (MD): NOAA - . [about 2 screens]. Available from: https://gml.noaa.gov/ccgg/trends/mlo.html

67. Bassirirad H. Kinetics of nutrient uptake by roots: Responses to global change. New Phytol. 2000;147(1): 155–169. doi: 10.1046/j.1469-8137.2000.00682.x

68. Alam SM. Nutrient uptake by plants under stress conditions. In: Pessarakli M, editor. Handbook of plant and crop stress. New York: Marcel Dekker; 1999. p. 285–313.

